# A modular transcriptional signature identifies phenotypic heterogeneity of human tuberculosis infection

**DOI:** 10.1101/216879

**Authors:** Akul Singhania, Raman Verma, Christine M. Graham, Jo Lee, Tran Trang, Matthew Richardson, Patrick Lecine, Philippe Leissner, Matthew P.R. Berry, Robert J. Wilkinson, Karine Kaiser, Marc Rodrigue, Gerrit Woltmann, Pranabashis Haldar, Anne O’Garra

**Author notes:** contributed equally (Akul Singhania, Raman Verma). co-supervised this work (Pranabashis Haldar, Anne O’Garra). Correspondence to AOG.

## Abstract

Whole blood transcriptional signatures distinguishing active tuberculosis patients from asymptomatic latently infected individuals exist. Consensus has not been achieved regarding the optimal reduced gene sets as diagnostic biomarkers that also achieve discrimination from other diseases. Here we show a blood transcriptional signature of active tuberculosis using RNA-Seq, confirming microarray results, that discriminates active tuberculosis from latently infected and healthy individuals, validating this signature in an independent cohort. Using an advanced modular approach, we utilise information from the entire transcriptome, which includes over-abundance of type I interferon-inducible genes and under-abundance of *IFNG* and *TBX21*, to develop a signature that discriminates active tuberculosis patients from latently infected individuals, or those with acute viral and bacterial infections. We suggest methods targeting gene selection across multiple discriminant modules can improve development of diagnostic biomarkers with improved performance. Finally, utilising the modular approach we demonstrate dynamic heterogeneity in a longitudinal study of recent tuberculosis contacts.

## Introduction

Tuberculosis (TB) is the leading cause of global mortality from an infectious disease. In 2016, there were 10.4 million incident and 6.3 million new cases of TB disease and 1.67 million deaths and its diagnosis is problematic^1^. Active pulmonary TB diagnosis requires culture of *Mycobacterium tuberculosis*, which may take up to 6 weeks^2^. Although the World Health Organisation^1^ endorsed GeneXpert MTB/RIF automated molecular test for *M. tuberculosis* results in rapid diagnosis^3^, this test still requires sputum which can be difficult to obtain. Difficulties in obtaining sputum lead to approximately 30% of patients in the USA and 50% of South African patients to be treated empirically^1,4^. However, clinical disease represents one end of a spectrum of infection states. An estimated one third of all individuals worldwide have been infected with the causative pathogen, *M. tuberculosis*, but the vast majority remain clinically asymptomatic with no radiological or microbiological evidence for active infection. This state is termed latent TB infection (LTBI) and conceptually denotes that *M. tuberculosis* persists within its host, while maintaining viability with the potential to replicate and cause symptomatic disease. Indeed, LTBI represents the primary reservoir for future incident TB, with 90% of all TB cases estimated to arise from reactivation of existing infection^1,5^. The risk of incident TB arising from existing LTBI is heterogeneous, poorly characterised and modifiable with anti-tuberculous treatment. Modelling studies indicate that effective TB prevention to reduce future TB incidence, requires policies directed at the identification and treatment of LTBI^6^. However, implementation of mass screening programmes for this purpose are severely constrained by the size of the target population. Transformative advances in diagnostic tools that can effectively help to stratify TB risk in the LTBI population are therefore implicit to the realisation of systematic screening.

The basis for LTBI heterogeneity rests with the limited scope of the tools we have available to identify the state. LTBI is inferred solely through evidence that immune sensitization has occurred, by the tuberculin skin test (TST) or the *M. tuberculosis* antigen-specific interferon-gamma (IFN-γ) release assay (IGRA). Although these tests are both sensitive and specific for identifying exposure, that has been associated with establishment of an adaptive immune response, neither distinguishes active from latent infection. Moreover, T-cell responses to mycobacterial antigens persist for several years after an infection has been treated, implying that these tests may not reliably inform the presence of viable organisms *in vivo*. For ‘true’ LTBI, in which the pathogen remains viable, it is envisaged that a dynamic equilibrium exists between the host immune response and the pathogen, with a shifting balance in favour of one or the other, influencing the future risk of TB reactivation^7^. A study using highly sensitive radiological imaging with combined Positron Emission Tomography and Computerised Tomography has reported evidence to support this dynamic state and demonstrated phenotypic imaging characteristics associated with the risk of developing TB among subjects with conventionally defined LTBI^8^. A proportion of these LTBI patients were identified with radiological features of subclinical active TB^8^, with a subgroup failing to respond to prophylactic LTBI treatment regimens. These observations support the view that injudicious use of LTBI chemoprophylaxis using presently available diagnostic tools for mass screening, risks promoting drug resistance in unrecognised active infection.

We have previously characterised an interferon (IFN) inducible transcriptional signature of 393 gene transcripts in whole blood that discriminates patients with active pulmonary TB (from high and low-incidence TB burden countries) from healthy individuals, patients with other chronic respiratory and systemic conditions, and the majority of patients with LTBI^9,10^. This TB signature revealed an unexpected dominance of type I IFN-inducible genes^9^ more frequently associated with viral infections^11^. We^12–14^ and others^13–23^ have since shown that elevated and sustained levels of type I IFN, result in an enhanced mycobacterial load and disease exacerbation in experimental models of TB. Similar findings of a blood signature in active TB patients have since been reported^24–30^, and our meta-analysis of 16 datasets, including many of these studies, identified 380 genes differentially abundant in active TB across all datasets^31^. However, there is a relative lack of concordance across studies that have reported a reduced and optimised diagnostic gene signature, although agreement exists for some of the pathways they represen^24–26,32,33^. While some genes overlap between the different reduced signatures, the overall composition of each reduced signature is unique, both in size and transcript profile. In this respect, we note that a consistent statistical approach to optimising gene selection has not been used across studies and where the approach was consistent, a different optimal reduced signature was reported for discriminating active TB from either LTBI and controls or other diseases^25^. Additionally, recent reports suggest these signatures do not effectively discriminate TB from other diseases such as pneumonia, lowering their value as stand-alone diagnostic test^29,30^.

We have previously observed and reported that 10 – 20% of subjects with IGRA positive LTBI in our studies had a transcriptional signature that overlapped with active TB patients and clustered with this group^9^. By definition, the transcriptional signature in this LTBI outlier group shares important similarities with the signature of active TB that requires further characterisation. Importantly, the biological significance of this statistical observation remains unclear. However, these observations support utilisation of a transcriptional approach to explore LTBI heterogeneity. In keeping with this, Zak et al.^33^ have recently reported evidence for a gene signature of TB several months in advance of clinical presentation with disease, among a cohort of South African adolescents, contained within the previously described TB signature^9^. This suggests that transcriptional signatures of TB in subjects with presumed LTBI may indicate either, a high risk of progression to active disease, or existing subclinical disease. However, interpretation of the study was limited by the confounding risk of new TB exposure in a high TB incidence setting, particularly to determine longitudinal changes in signature expression and dynamic heterogeneity of the host immune response. Analysis focussed on the subgroup with IGRA defines LTBI, despite a proportion of prospective TB cases developing in subjects that were IGRA negative at baseline, suggesting either new TB exposure during prospective observation in a high TB incidence setting and/or that the IGRA test did not reliably inform underlying LTBI. In this context, studies evaluating the diagnostic performance of IGRAs in microbiologically confirmed active TB report an overall sensitivity of approximately 85%^<u>34</u>^, implying that a proportion of *M. tuberculosis* infections may be missed using this test alone.

To address some of these questions, here we: (i) validate microarray findings by RNA-sequencing in our published^9^ cohorts and a new cohort of TB; (ii) evaluate current TB gene signatures from the literature against TB and other infections; (iii) develop and test a modular TB signature in multiple TB cohorts and other diseases; (iv) develop a reduced TB-specific gene signature from modules of TB and test against viral infections and other diseases; (v) evaluate the transcriptional profile of LTBI-Outliers and compare with active TB; (vi) evaluate the blood transcriptional signature at baseline and longitudinally in TB contacts who develop TB against those TB contacts who remain healthy, in a low TB incidence setting (**Figure 1; Supplementary Figure 1**). As a proof of principle, our reduced TB-specific gene set developed from the modular signature, not only distinguishes active TB and LTBI but additionally does not detect viral and bacterial infections. We identify immunological heterogeneity of LTBI, with a percentage of individuals showing a transcriptional signature of active TB, which only develop longitudinally in a small proportion of the recent close contacts of TB.

**Figure 1.**
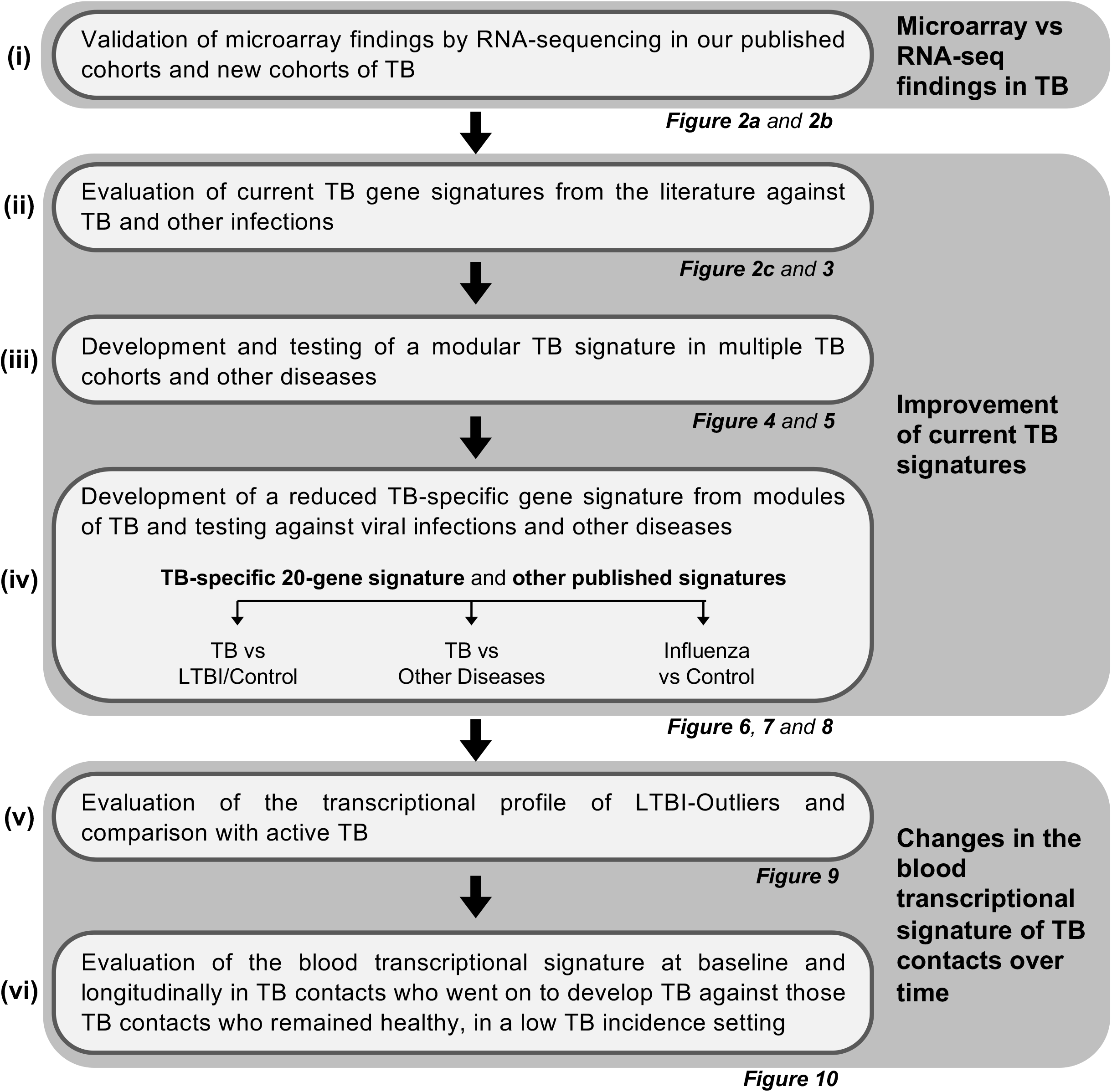
The objectives of this study. An overview of the analysis undertaken in the study. Figures associated with each objective are stated below the box.

## Results

### RNA-seq recapitulates the microarray TB gene signature

We validated our microarray-derived blood 393-transcript signature^9^ in patients with active TB using RNA-Seq in the Berry London and South Africa cohorts showing identical clustering of active TB and LTBI cases (**Supplementary Figure 2a** and **2b**). A 373-gene signature was then independently re-derived from the Berry London RNA-Seq data (Supplementary **Figure 2c; Supplementary Data 1; Figure 2a**) and validated in the Berry South Africa cohorts (**Figure 2a**) and a new Leicester cohort (**Supplementary Table 1; Figure 2b**). Consistent with our previous microarray signature, the RNA-Seq signature was absent in the majority of individuals with LTBI and healthy controls, and identified with perfect agreement the LTBI subjects that cluster with active TB, henceforth referred to as LTBI outliers, in both Berry cohorts (**Supplementary Figure 2b and 2d**). A similar proportion of outliers were also observed in the Leicester cohort (**Figure 2b; Supplementary Figure 2e**). There was great similarity in the composition of the microarray and RNA-Seq based signatures, with over-abundance of IFN-inducible genes and under-abundance of B- and T-cell genes as previously reported^9^. This was supported by an *in silico* cellular deconvolution analysis of the RNA-Seq that showed diminished percentages of CD4, CD8 and B cells in the blood of active TB patients, and an increase in monocytes/macrophages and neutrophils (**Supplementary Figure 3**), in keeping with our previous findings using flow cytometry^9^.

**Figure 2.**
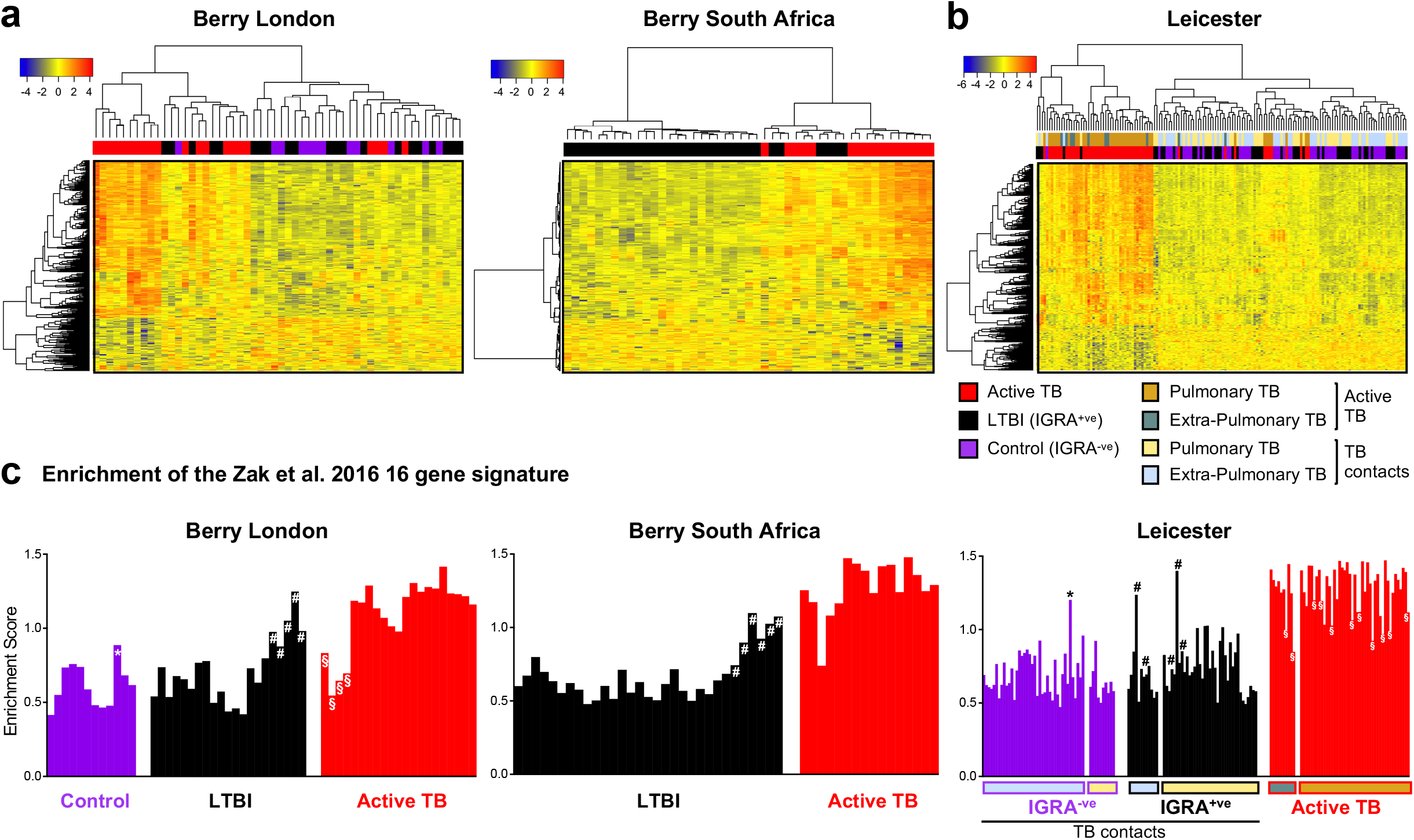
Whole-blood transcriptional gene signatures in TB. **a** Heatmaps depicting unsupervised hierarchical clustering of active TB (red), LTBI (black) and control samples (purple) using a 373-gene signature derived using the Berry London cohort, tested in the Berry South Africa cohort, and (**b**) validated in an independent cohort from Leicester. Gene expression values were averaged and scaled across the row to indicate the number of standard deviations above (red) or below (blue) the mean, denoted as row Z-score. c Bar graphs depicting enrichment scores derived on a single sample basis using ssGSEA in the Berry London, Berry South Africa and Leicester cohorts using the 16-gene signature from Zak et al. 2016^33^. Purple, black and red bars represent control, LTBI and active TB samples, respectively, and * (control outliers), # (LTBI outliers) and § (active TB outliers) represent outlier samples identified by hierarchical clustering.

### Published TB gene signatures identify acute viral infections

Applying the published 16-gene signature of Zak et al.^33^ to the Berry and Leicester TB cohorts, single sample Gene Set Enrichment Analysis^35^ (ssGSEA) across all three cohorts, demonstrated high enrichment of the Zak et al. signature^33^ in active TB and a low enrichment in healthy controls and the majority of LTBI patients (**Figure 2c**). We observed higher enrichment scores in the LTBI outlier groups (**Figure 3a and 3b**) of all three cohorts that overlapped with scores observed in active TB cohorts (**Figure 2c**). Higher enrichment scores were also noted in a small proportion of IGRA^−ve^ individuals recruited as healthy controls (**Figure 2c**). There was comparable discrimination in enrichment scores between TB and LTBI in the Zak et al.^33^ 16-gene signature, and the Kaforou et al., 27 and 44-gene signatures^25^ (**Figure 3a**). Only the Kaforou 44-gene signature was developed to discriminate between active TB and other diseases (including infectious meningitis, pneumonia, gastric diseases and malignancies)^25^, rather than LTBI.

**Figure 3.**
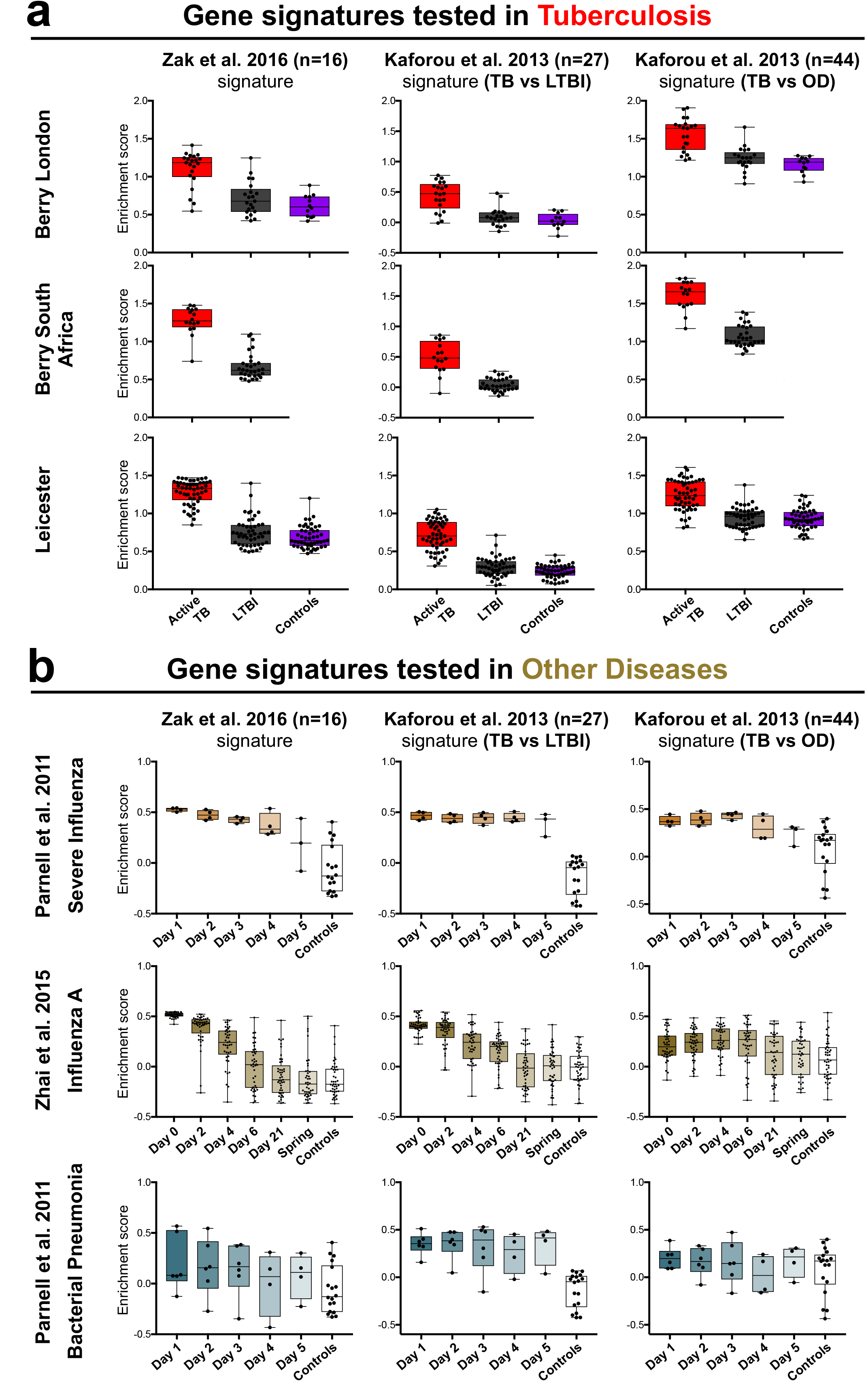
Enrichment of published reduced TB gene signatures in TB, and other viral and bacterial infections. **a** Box plots depicting enrichment scores derived on a single sample basis using ssGSEA, using the 16-gene signature from Zak et al. 2016^33^, and the 27-gene (TB vs. LTBI) and 44-gene (TB vs. other diseases (OD)) signatures from Kaforou et al. 2013^25^ in tuberculosis datasets (Berry London, Berry South Africa and Leicester), and (**b**) in datasets of other infections - severe influenza from Parnell et al.^36^, Influenza A from Zhai et al.^37^, and bacterial pneumonia from Parnell et al.^36^. The box represents 25th to 75th percentile, with a line inside the box indicating the median, and the whiskers represent the minimum to the maximum points in the data.

The composition of all three signatures^25,33^ is dominated (≥50% of the signature) by IFN-inducible genes (**Supplementary Table 2**), raising the possibility that they are not TB specific but may also be expressed in acute viral infections. We therefore evaluated enrichment of these signatures in two independent published datasets of influenza infection from Parnell et al.^36^ and Zhai et al.^37^ (**Supplementary Table 3; Figure 3b**). Subjects with influenza at baseline showed a high enrichment score for the three TB signatures as compared with healthy controls, which diminished with time, in keeping with recovery (**Figure 3b**). In contrast, enrichment scores for the three signatures, demonstrated heterogeneity in patients diagnosed with bacterial pneumonia from the Parnell study^36^, with little change over 5 days and poor discrimination from controls, consistent with our previous findings for this group^9,10^ (**Figure 3b**).

### A modular signature discriminates TB from other diseases

A limitation of the gene reduction methodologies^25,33^ used to date has been the prioritisation of the most discriminant genes, with little consideration to the correlation between the selected genes in this iterative process. Although non-selective and lacking subjective bias, this approach favours selection of a highly correlated gene set with a narrow immunological focus. In this context, limited diversity risks loss of specificity, with an increased likelihood of overlap between multiple pathologies and responses to different infections for a specific immune pathway. We therefore hypothesised that methodologies which incorporate information from the entire transcriptome may better inform development of a unique biosignature for TB. Weighted gene co-expression network analysis^38^ (WGCNA) is a well validated clustering technique for reducing high dimensional data into modules that preserve intrinsic relationships between variables within a network structure. When applied to the blood transcriptome, modules of co-ordinately expressed genes with a coherent functional relationship are generated. The complete transcriptome is thus expressed as a signature defined by the relative perturbation of individual modules.

We applied WGCNA analysis to the blood transcriptional data from our Berry and Leicester TB cohorts, those TB cohorts published by Zak and Kaforou^25,33^, and to several others that included sample sets of other viral and bacterial infections^36,37,39,40^, together with our previous cohorts of sarcoidosis and lung cancer^10^ as conditions that may mimic TB, all compared against their healthy controls (**Figure 4; Supplementary Table 3** (Information of published cohorts), **Supplementary Data 2** (genes in each module) and **Supplementary Table 4** (module annotation)). The modular signature for active TB was qualitatively consistent across all the TB cohorts and absent in LTBI. The IFN-modules (lightgreen and yellow) were over-abundant in TB (**Figure 4a**) as we have previously published^9,10^ and also in acute influenza infection, but absent in bacterial infection^9,10^ (**Figure 4b**). However, we observed clear differences between TB and both influenza and other bacterial infections in the pattern of specific perturbation of other modules, including under-abundance of gene expression in the T-cell (blue and cyan) and B-cell (midnightblue) modules (**Figure 4**) for TB. On the other hand, we observed over-abundance of genes in the Cell Proliferation/Metabolism (darkturquoise) module and under-abundance of genes associated with Haematopoeisis (pink) in severe influenza but not in TB (**Figure 4**). In this context, the classical approaches of gene signature reduction algorithms^41–43^ used by Kaforou et al. to distinguish TB from LTBI or TB from other diseases^25^, and Zak et al. to distinguish TB from LTBI, risk of progression^33^ are notable for formulating gene signatures that we show here map predominantly to the yellow module (Interferon/complement/myeloid), with many of these genes also over-abundant in both influenza cohorts (**Supplementary Figure 4 and 5**).

**Figure 4.**
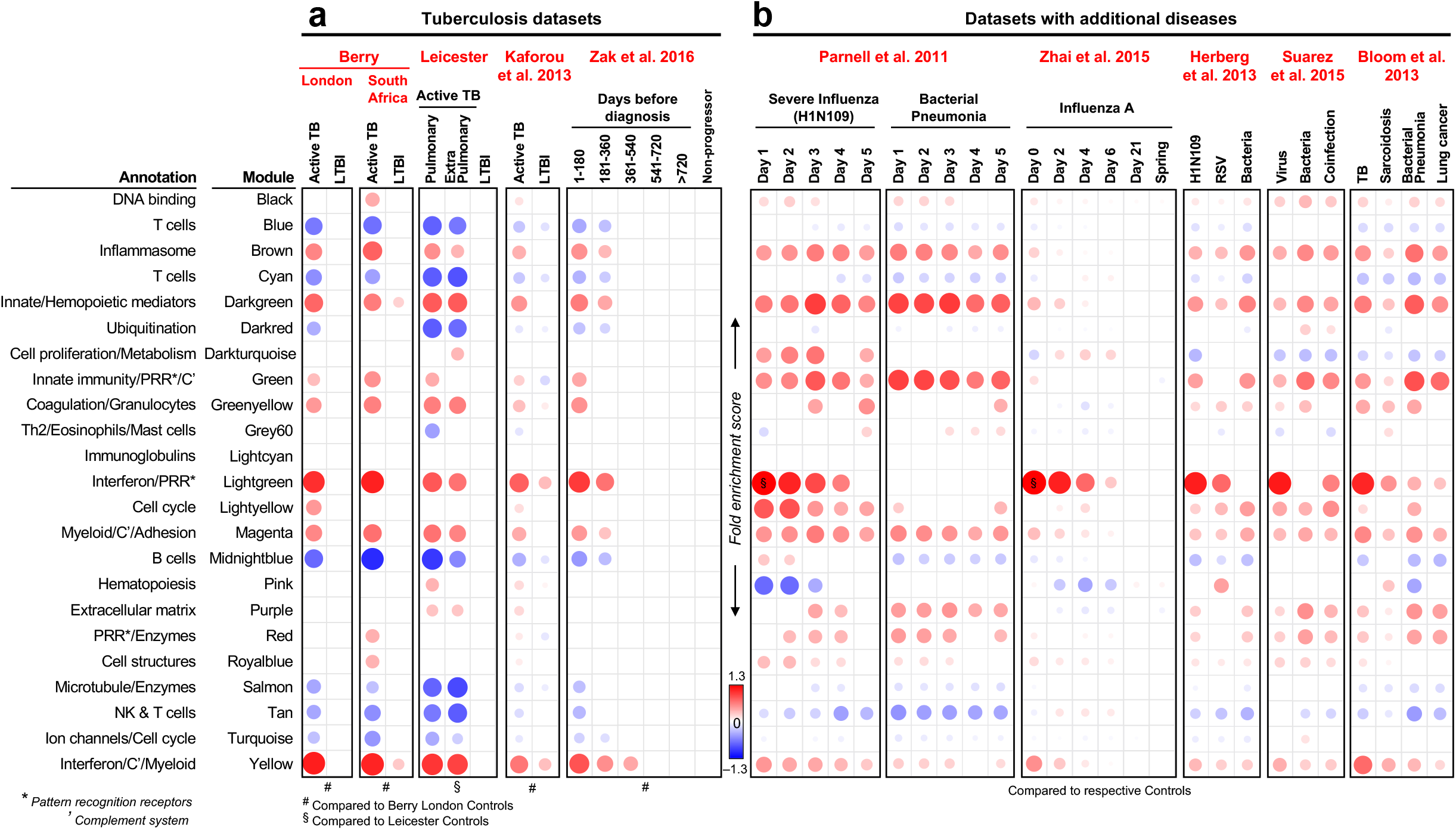
Modular transcriptional signatures of TB and other diseases. **a** Twenty-three modules of co-expressed genes derived using WGCNA from Combined Berry dataset (London & South Africa) and tested in other TB datasets, and (b) datasets with additional diseases. Fold enrichment scores derived using QuSAGE are depicted, with red and blue indicating modules over- or under-expressed compared to controls. Colour intensity and size represent the degree of enrichment compared to controls. Only modules with fold enrichment scores with FDR p-value < 0.05 were considered significant and depicted here. §, fold enrichment scores in the lightgreen module greater than the maximum score depicted on the scale (i.e. >1.3) in severe influenza (Parnell et al.^36^, score: 1.55) and influenza A (Zhai et al.^37^, score: 1.97).

### A reduced TB-specific signature from modular gene expression

Interrogating the whole gene-set of the yellow module in TB, influenza and bacterial infection, we observed a subset of genes expressed specifically in TB (**Figure 5, orange squares**). Similarly, other genes were specifically expressed in influenza. Thus, although modular expression of the yellow module is comparable between TB and influenza, gene subsets within the module exhibit differential expression between the two conditions. This provides scope to select genes from this dominant module that can be used to develop a TB signature, while retaining discriminant value from viral infection. Using this rationale, as a proof of principle, we identified and extracted 303 unique gene candidates in the Berry London TB dataset that were selectively perturbed in TB, but not in any confounding viral infections, from all modules that contributed to and exhibited consistency across the TB datasets that we analysed (**Figure 4; Supplementary Figure 6a; Supplementary Figure 6b**). Using this gene set, we developed a reduced gene signature to distinguish active TB from LTBI. We applied the Boruta algorithm^42^ based on random forest to this set of genes, yielding 61 genes (**Supplementary Figure 6c**) that was further reduced by selecting the top 20 genes, ranked according to GINI score using Random Forest (**Supplementary Figure 6d**). Our 20-gene signature (**Figure 6a**) included genes from six different modules (**Supplementary Figure 6d**), representing both over-abundance and under-abundance in TB. Using a modified Disease Risk Score (See Methods), we identified powerful discrimination between active TB and LTBI/controls in Berry London & South Africa and Leicester cohorts (**Figure 6b**). In contrast, the signature identified no difference between influenza and controls or between bacterial pneumonia and controls at any time-point across five days (**Figure 6c**). Our 20-gene signature also discriminated active Tb from LTBI and controls, in three additional published cohorts, similarly to the 44-gene signature described by Kaforou et al.,^25^ (**Figure 7a**). Both our 20-gene signature and the 44-gene signature of Kaforou also discriminated active TB from other diseases, albeit to a lower extent (**Figure 7b**). In keeping with this, our 20-gene signature, and the signatures published by Zak et al.^33^, Kaforou et al.^25^, Roe et al.^28^, Sweeney et al.^44^, Maertzdorf et al.^45^, distinguished active TB and LTBI with high specificity and sensitivity (**Figure 8a**). Whereas our 20-gene signature did not discriminate influenza from controls, all other signatures demonstrated excellent discrimination between influenza from controls, comparable with their performance for TB (**Figure 8b**), indicating that influenza and other types of viral infections may inadvertently be detected and confound the TB diagnosis.

**Figure 5.**
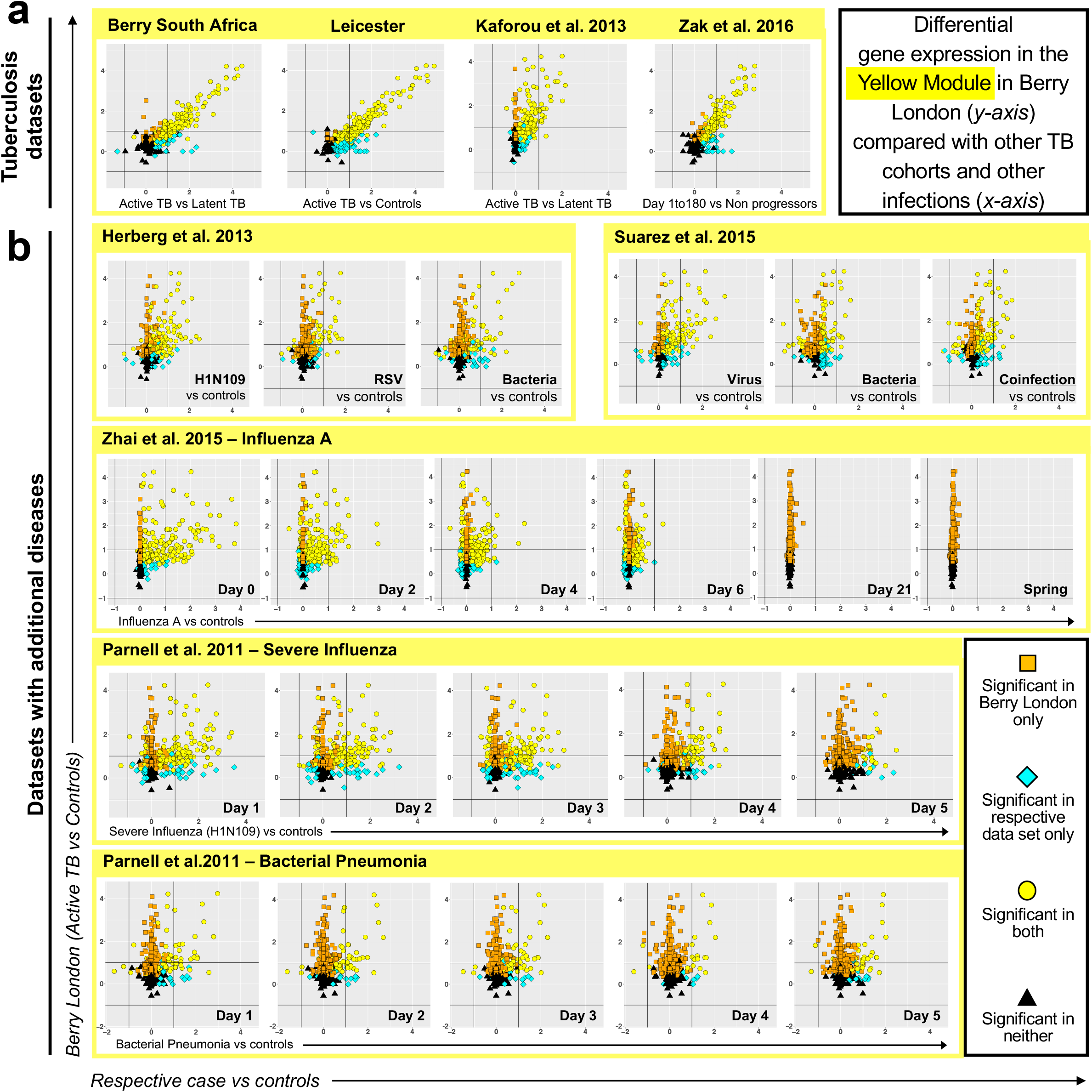
Gene expression in the yellow module in TB compared to other viral and bacterial infections. **a** Log_2_ fold changes for genes in the yellow module from Berry London cohort (active TB vs. controls; *y-axis*) compared to log_2_ fold changes in other datasets (respective cases vs. controls; *x-axis)* in TB, and (**b**) other infections (Herberg et al. 2013^39^, Suarez et al. 2015^40^, time-course data from Zhai et al. 2015^37^ (influenza A) and Parnell et al. 2011^36^ (severe influenza and bacterial pneumonia)). Shapes and colours represent significantly differentially expressed genes (FDR p-value < 0.05) in either Berry London only (orange squares), respective dataset only (cyan diamonds), both dataset (yellow circles) or significant in neither (black triangles).

**Figure 6.**
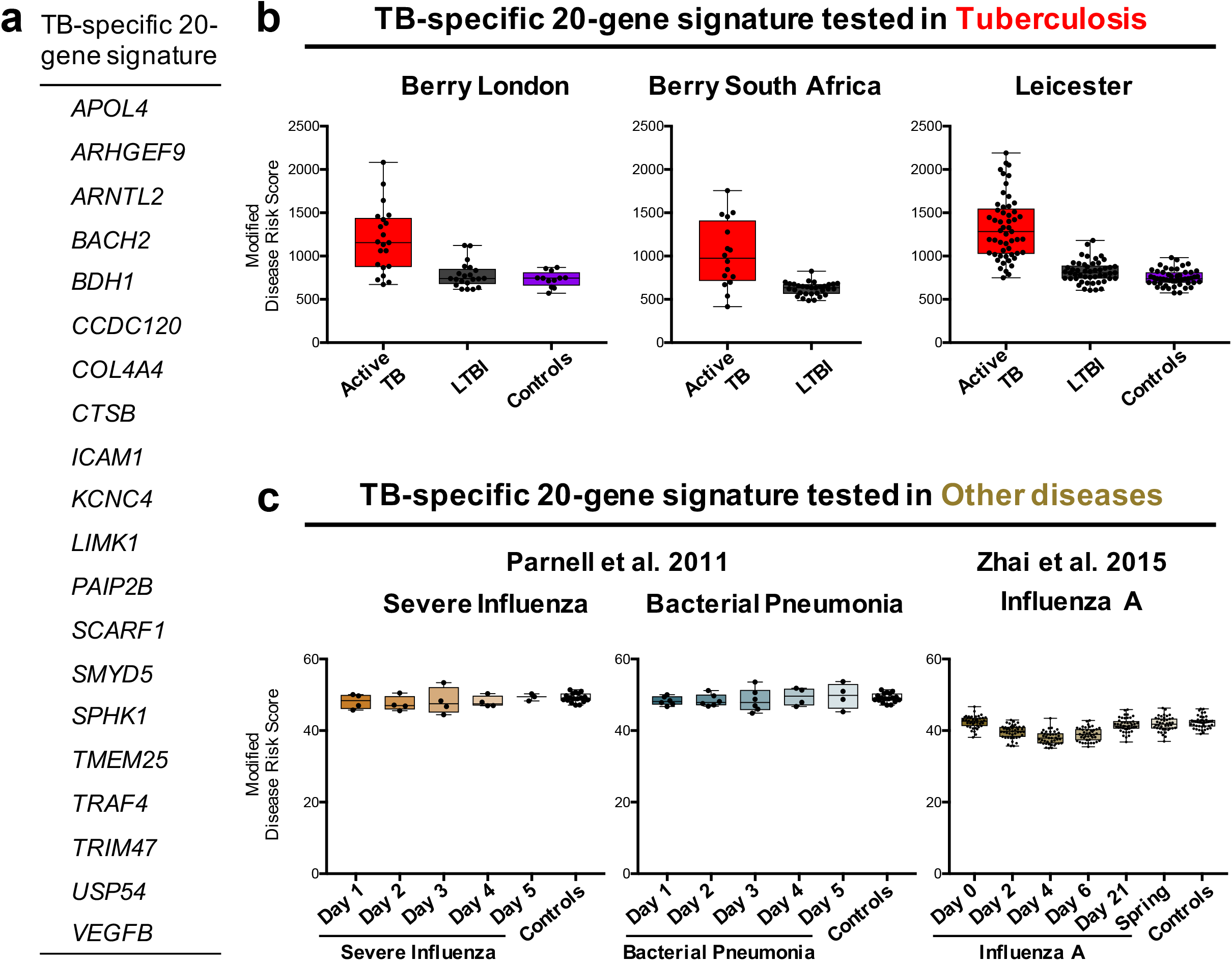
Whole-blood TB-specific 20-gene signature tested in TB and other infections. **a** A reduced 20-gene signature of TB derived from the TB modular signature using genes significantly differentially expressed in Berry London cohort only, and not in other flu datasets (**Supplementary Figure 6 5**). **b** Box plots depicting the modified Disease Risk Scores derived using the TB-specific 20-gene signature in TB datasets, and in (**c**) datasets of other infections. The box represents 25th to 75th percentile, with a line inside the box indicating the median, and the whiskers represent the minimum to the maximum points in the data.

**Figure 7.**
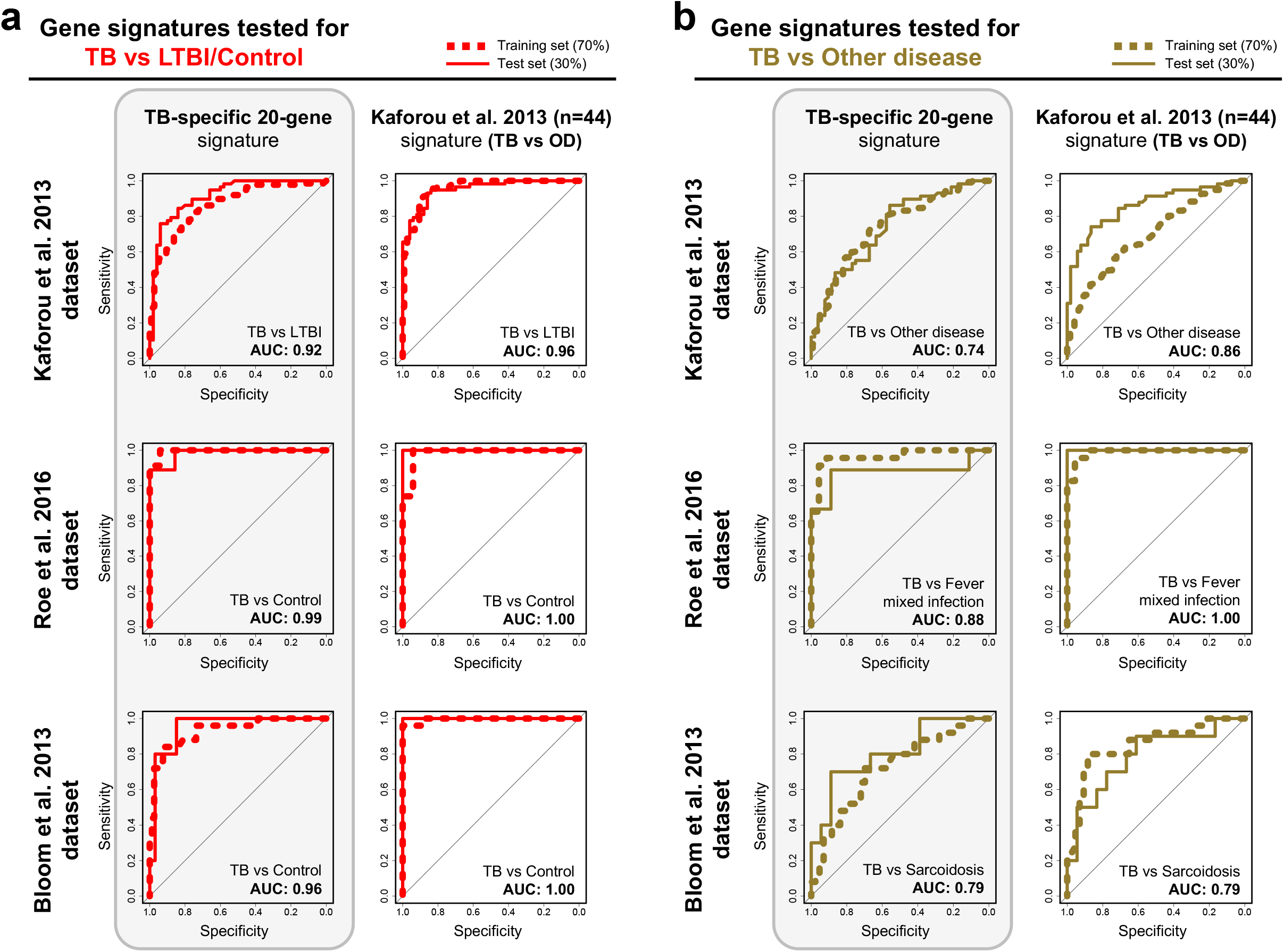
Comparison of our TB-specific 20-gene signature with Kafourou’s in distinguishing TB and other diseases. **a** Receiver operating characteristic (ROC) curves depicting the predictive potential of the TB-specific 20-gene signature and the 44-gene (TB vs. other diseases (OD)) signature from Kaforou et al. 2013^25^ in classifying a sample as TB or LTBI/Control, or (**b**) in classifying a sample as TB or other disease, in datasets from Kaforou et al. 2013^25^, Roe et al. 2016^28^ and Bloom et al. 2013^10^. Area under the curve (AUC) is shown for each ROC curve.

**Figure 8.**
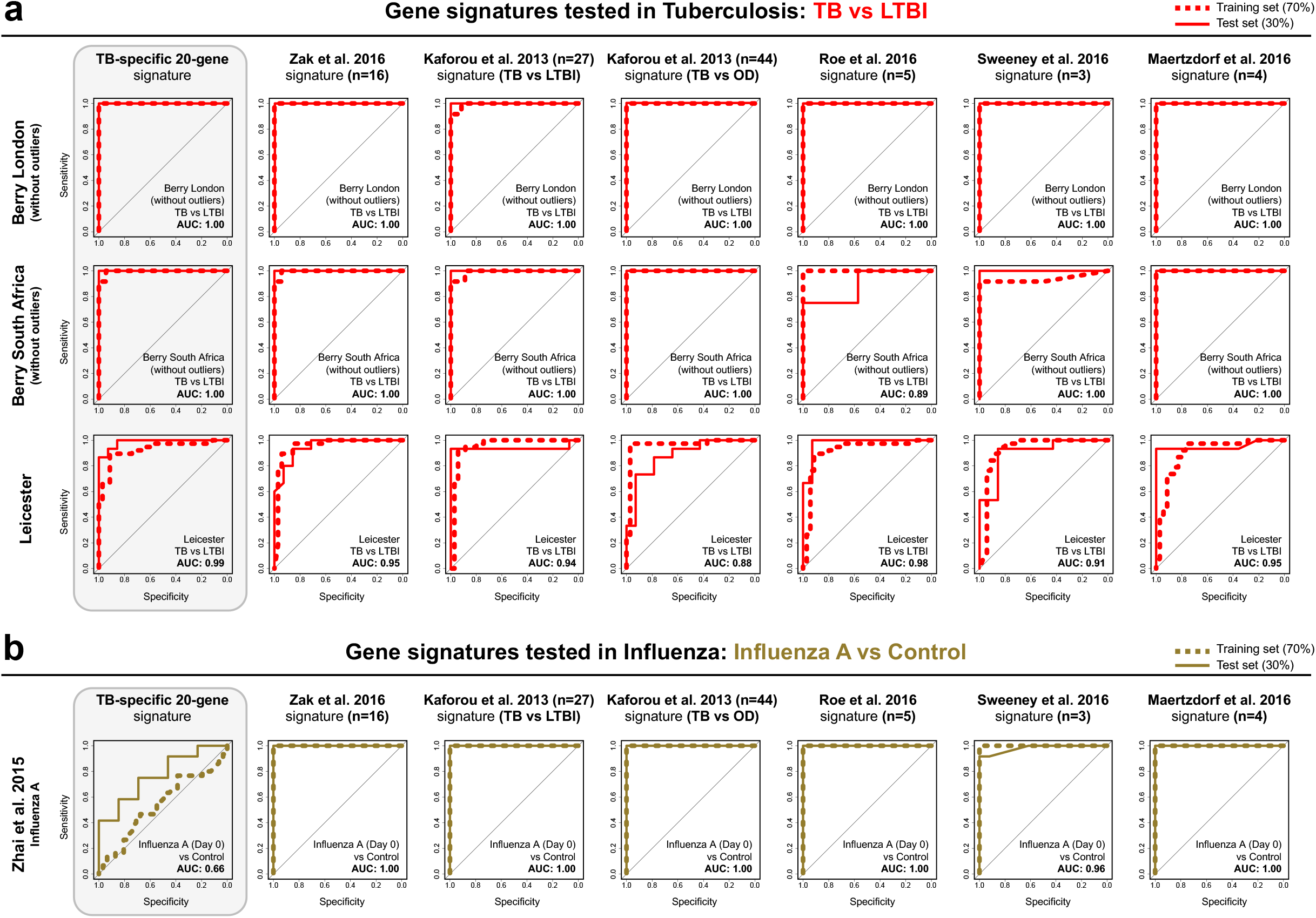
Comparison of our TB-specific 20-gene signature and others in distinguishing TB and influenza. **a** Receiver operating characteristic curves (ROC) depicting the predictive potential of the TB-specific 20-gene signature and the other published gene signatures in classifying a sample as TB or LTBI in the Berry London, Berry South Africa and Leicester cohorts, and (**b**) in classifying a sample as influenza A or control in the Zhai et al. 2015^37^ dataset. Area under the curve (AUC) is shown for each ROC curve.

### The modular signature of LTBI outliers resembles that of TB

We have previously reported evidence for a small proportion of LTBI subjects that clustered with active TB using our 393-transcript signature^9^ that we refer to as an LTBI outlier group. This group was reproduced using RNA-Seq in the Berry cohorts (10.9%) and a similar proportion were also identified in our new Leicester cohort (10%) (**Figure 2**). To compare and contrast the signature of this group with active TB and the majority of LTBI resembling healthy controls (**Supplementary Figure 2d** and **2e**), we specifically examined the WGCNA modular signature in LTBI outliers using the combined Berry London and South Africa datasets and Leicester datasets respectively, compared with healthy controls (**Figure 9a**). The modular signature of LTBI outliers in both datasets showed over-abundance of the lightgreen (IFN/Pattern recognition receptors) and yellow (IFN/Complement/myeloid) modules as seen in active TB (**Figure 9a and 9b**). This is entirely in keeping with our earlier finding (**Figure 2c and 3a**) that gene enrichment scores using the published signatures^25,33^, all of which are comprised primarily of genes from the yellow module (**Supplementary Figures 4 and 5**), were consistently higher in LTBI outliers. Of note, these gene-sets were not present in the lightgreen (IFN/Pattern recognition receptors). In addition to overabundance of the IFN modules, the LTBI outlier group of the Leicester dataset showed changes in other modules also perturbed in active TB, suggesting a host response that is evolving towards the phenotype typically observed in active TB (**Figure 9a**). Of particular interest was the observation of under-abundance in the tan module (Th1 and NK cells) that is associated with IFN-γ expression, a cytokine required for protection against TB^16,46–52^. Under-abundance of this module was a consistent finding across all the TB datasets that we analysed (**Figure 4, Figure 9**).

**Figure 9.**
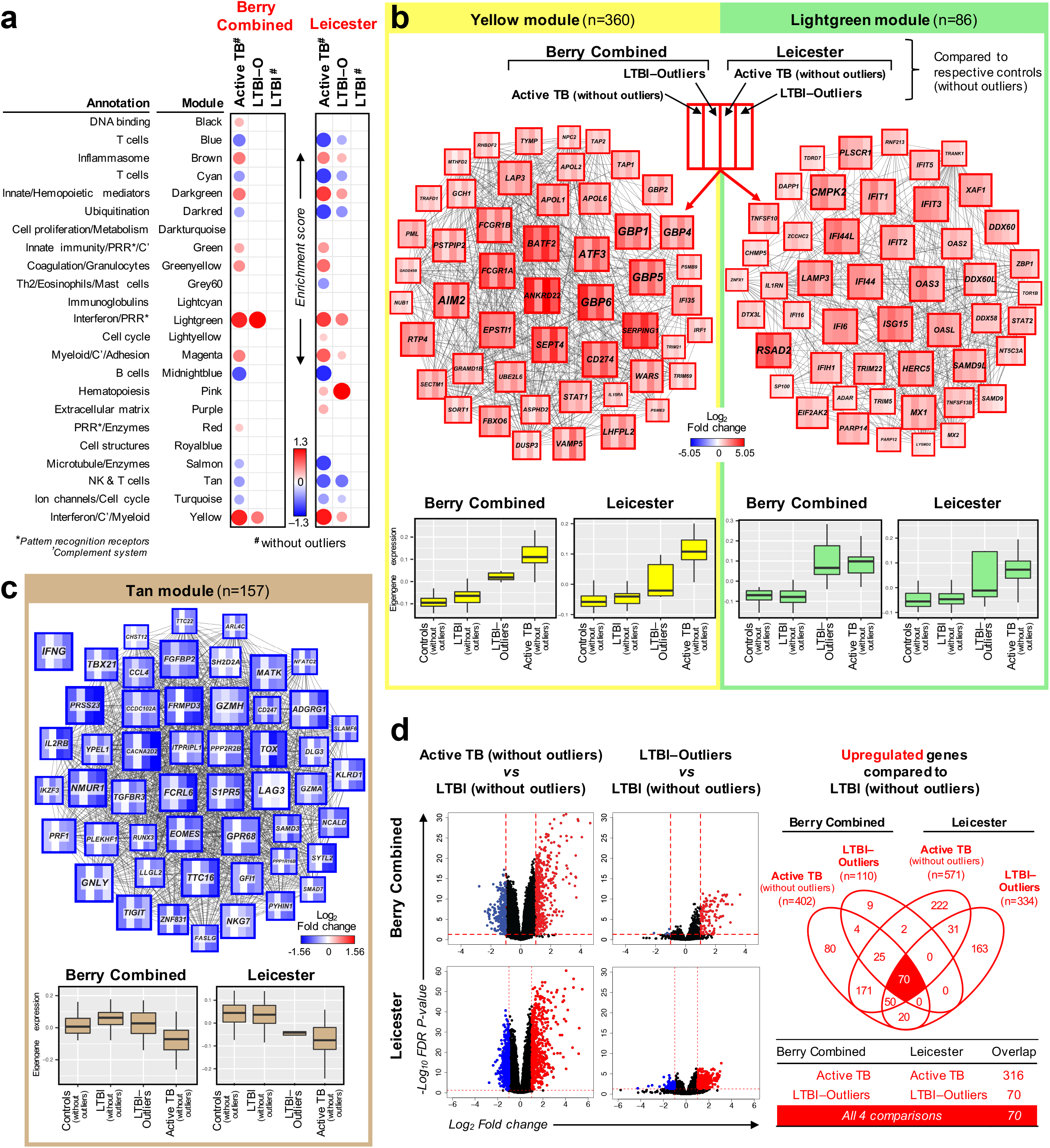
Blood transcriptional profile of LTBI outliers compared with active TB. **a** Modules of co-expressed genes tested in LTBI-Outliers from the Combined Berry and Leicester cohorts. Fold enrichment scores derived using QuSAGE are depicted, with red and blue indicating modules over- or under-expressed compared to controls. Colour intensity and size represent the degree of enrichment compared to controls. Only modules with fold enrichment scores with FDR p-value < 0.05 were considered significant and depicted here. **b** Gene networks depicting the top 50 ‘hub’ genes, i.e. genes with high intramodular connectivity, for the yellow, lightgreen and (**c**) tan modules. Each gene is represented as a square node with edges representing correlation between the gene expression profiles of the two respective genes (minimum Pearson correlation of 0.75). A key describing the four different partitions within each square node is shown, with each partition representing log2 fold changes for active TB (without outliers) and LTBI-Outliers from the Berry Combined and Leicester cohorts, compared to respective controls (without outliers). Red and blue represent up- and down-regulated genes, respectively. In the tan module, the expression for IFNG is also shown, although it was not one of the top 50 hub genes within that module. Boxplots depicting the module eigengene expression, i.e. the first principal component for all genes within the module, are shown below each gene network. **d** Volcano plots depicting differentially expressed genes for active TB (without outliers) and LTBI-Outliers in the Berry Combined and Leicester cohorts, compared to respective LTBI (without outliers). Significantly differentially expressed genes (log2 fold change >1 or <−1, and FDR p-value < 0.05) are represented as red (upregulated) or blue (downregulated) dots, along with a Venn diagram and table summarising overlaps between these different comparisons.

We performed differential gene expression analysis between active TB, LTBI outliers, and LTBI with outliers removed, and identified a set of 70 genes that was consistently upregulated in active TB and LTBI outliers compared to LTBI (without outliers) in both the Berry and Leicester datasets (**Figure 9d; Supplementary Data 3**), which were enriched for the IFN signalling pathway and innate immunity.

### Dynamic transcriptional heterogeneity in recent TB contacts

Longitudinal RNA-Seq was performed in a subset of our Leicester cohort (**Methods; Figure 10a**) that included 15 IGRA^−ve^ contacts, 16 IGRA^+ve^ contacts, both of whom remained healthy, and 9 subjects recruited as contacts that were subsequently diagnosed with microbiologically confirmed TB during prospective observation (**Figure 10a; Supplementary Table 5**). Five contacts (4 IGRA^+ve^ and 1 IGRA^−ve^) identified as outliers at baseline sequencing (**Figure 2b**) were included.

**Figure 10.**
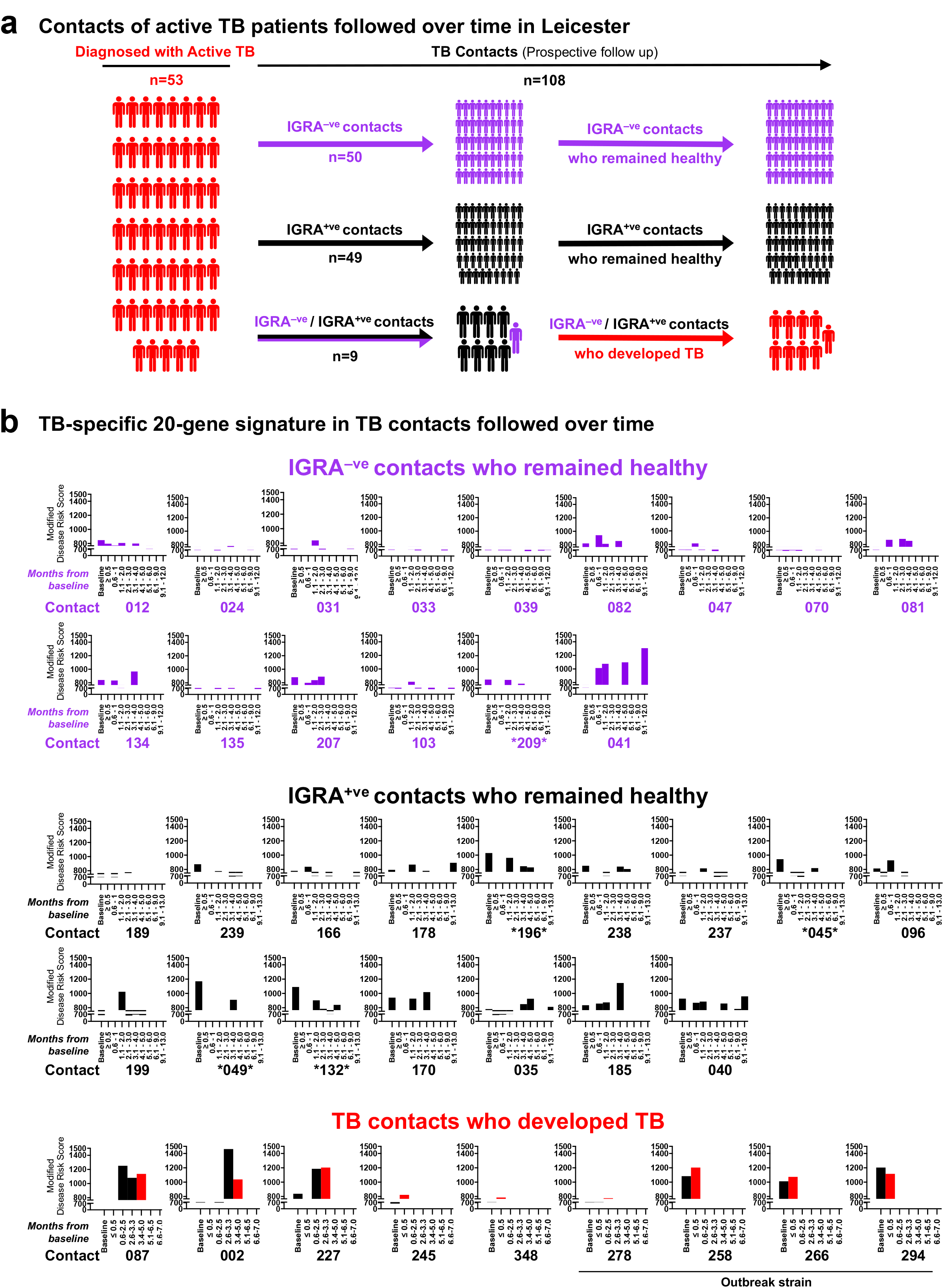
Blood transcriptional profile of TB contacts followed over time. **a** Schematic diagram representing active TB patients from the Leicester cohort, and their contacts followed over time. Purple, black and red represent IGRA^−ve^ (controls), IGRA^+ve^ (LTBI) and active TB patients, respectively. **b** Bar plots depicting the modified Disease Risk Scores using the TB-specific 20-gene signature, in TB contacts who remained IGRA^−ve^ and did not develop TB (n=15), TB contacts who remained IGRA^+ve^ and did not develop TB (n=16), and TB contacts who developed TB during the study (n=9). For TB contacts who developed TB during the study, the time point when the contact was diagnosed with active TB in the clinic is represented by a red bar. Baseline in the barplot is set at 766.64, average of all Baseline time-point modified Disease Risk Scores from all IGRA^−ve^ contacts (n=15).

In contrast with other studies, the control population of our Leicester cohort comprised subjects that were IGRA^−ve^ contacts of TB. This is a group in which recent exposure to active TB is documented, placing them at higher risk of recently acquired infection. Our rationale for this approach was to evaluate whether blood transcriptional data may identify LTBI that is not detected using IGRA. The observations that: firstly, the Leicester control group had greater overlap in enrichment scores with the IGRA^+ve^ LTBI group using the Zak and Kaforou signatures, compared with the Berry London cohort (**Figure 2c and 3a**); and secondly, one subject from this group was identified as an outlier, together suggest that IGRA testing alone may miss some *M. tuberculosis* infection. We therefore elected to define our TB contacts henceforth as IGRA^+ve^ or IGRA^−ve^ with no deterministic reference to LTBI.

The modular signatures of both IGRA^−ve^ and IGRA^+ve^ contacts qualitatively demonstrated considerable between-subject heterogeneity and some within-subject variability; a comparison between the groups suggested more transcriptional activity, in the form of a higher frequency and greater breadth of modules exhibiting overabundance and under-abundance within the IGRA^+ve^ group (**Supplementary Figure 7**). For the cohort that developed TB after recruitment to the study (**Supplementary Figure 7**), we stratified subjects on the basis of their longitudinal clinical course as true progressors (no evidence of TB at baseline, with features developing during observation); subclinical TB (objective evidence of pathology, usually as radiological change, in the absence of reported symptoms); and active TB (symptoms at baseline with either radiological or microbiological evidence for TB subsequently identified) (**Supplementary Table 5**). This stratification was performed to better understand the dynamic relationship between the modular signature and onset of TB.

To quantitatively evaluate the modular signatures in each group for their proximity to TB, we applied the modified disease risk score for our 20-gene signature (**Figure 10b**). Higher risk scores were generally observed in the IGRA^+ve^, compared with the IGRA^−ve^ cohort, although there was considerable variability and overlap. Longitudinal observations suggest relative stability of the risk score in the majority of both IGRA^+ve^ and IGRA^−ve^ subjects that were examined. In contrast, 6 of the 9 subjects that were diagnosed with TB demonstrated high baseline modified disease risk scores that tended to increase further, prior to diagnosis of active TB. In the other 3 contacts (Subjects 245, 348 and 278) the modified disease risk score remained low at all time-points, before and at the time of TB diagnosis (**Figure 10b**).

Baseline scores demonstrated clustering near baseline for IGRA^−ve^ subjects (**Supplementary Figure 8; Figure 10b**). In contrast, both the IGRA^+ve^ group, and the group that developed TB exhibited a higher risk score (**Figure 10b; Supplementary Figure 8**). Subjects identified as outliers (**Figure 2b**), marked with an asterisk* (**Figure 10b**), had higher TB scores than the majority of LTBI subjects that were not outliers (**Figure 10b**). However, some discordance between clustering outcomes (**Figure 2b**) and the TB score was observed (**Figure 10b**), with two subjects that were not outliers having TB scores within the outlier range (subject 185 and 040). Furthermore, the IGRA^−ve^ subject categorised as an outlier (Subject 209) had a very low TB score (**Figure 10b**). Overall, the longitudinal within-subject expression of the 20-gene TB signature in both the IGRA^+ve^ and IGRA^−ve^ cohorts could be categorised into three groups: i. Subjects that did not express the signature at any time-point (10 of 15 IGRA^−ve^ subjects and 6 of 16 IGRA^+ve^ subjects); ii. Subjects that transiently expressed the signature, albeit generally to a low extent, in the first three to four months (4 of 15 IGRA^−ve^ subjects and 8 of 16 IGRA^+ve^ subjects; iii) Subjects that had or developed a persistent TB signature at and beyond 4 months (1 of 15 IGRA^−ve^ subjects and 2 of 16 IGRA^+ve^ subjects)(**Figure 10b**). We did not observe subjects developing the signature *de novo* after 3 months. Using the 16-gene signature of Zak et al.^33^, we report similar findings (**Supplementary Figure 9**). However, this 16-gene signature showed scores for additional IGRA^−ve^ and IGRA^+ve^ individuals (**Supplementary Figure 9, marked with red arrow**), possibly resulting from intercurrent infections.

In the cohort that developed TB, 5 of the 9 subjects demonstrated high baseline modified disease risk scores (**Figure 10b**). In 6 of the 9 subjects a moderate to high TB score was observed at the visit prior to TB diagnosis. For the remaining 3 subjects (Subject 245, Subject 278 and Subject 348), a 20-gene signature of TB was not expressed. For Subject 245 an explanation may be that this patient received antibiotics for bacterial pneumonia which are known to have immunosuppressive effects that may have affected expression of the immune signature. For Subjects 278 and 348 we have not identified potential confounding factors for this observation. Subjects categorised as true progressors exhibited a dynamic modular signature, with increasing TB scores at all visit time-points within 2 months of diagnosis.

## Discussion

We have recapitulated a blood transcriptional signature of active TB using RNA-Seq, previously reported by microarray^9,25,26,31,53^ that discriminates active TB from LTBI and healthy individuals, and is largely characterised by an over-abundance of IFN-inducible genes and an under-abundance of B and T cell genes. We show that an advanced modular approach, rather than a traditionally derived reduced gene set, is robust in discriminating active TB patients from individuals with LTBI whilst additionally not detecting acute viral and bacterial infections. Our findings highlight the need to consider additional approaches to develop transcriptional biomarkers of the highest sensitivity and specificity for distinguishing active TB from LTBI and other diseases. Using this modular approach and our reduced gene set, we also demonstrate heterogeneity of LTBI in a prospective study of contacts of patients with active TB.

RNA-Seq^33^ has now replaced microarray^9,10,24–27,31,32,53–55^ for transcriptional studies and the existing literature is limited by uncertainty regarding the equivalence of RNA-Seq and microarray. In this study, we repeated analysis of our previous Berry et al.^9^ cohorts using RNA-Seq and provide reassurance that RNA-Seq recapitulates outcomes derived using microarray. The vast majority of genes in our RNA-Seq derived 373-gene signature also comprised our original 393-gene transcript signature. Furthermore, there was equivalence in allocation of subjects to clusters, including those with LTBI that clustered with active TB and thus referred to as outliers. This is of considerable translational significance as no field applicable test is likely to be based on RNA-seq or microarray analysis.

In transcriptomic studies of disease there has been focus on deriving reduced gene signatures to develop clinical diagnostics, with inconsistencies in both deriving and defining the optimal reduced gene signature. Studies defining signatures distinguishing active TB and LTBI^25,26,28,33,44,56^ are illustrative of this issue. We evaluated the diagnostic performance of some of the published reduced gene signatures^25,28,33,44,45^ on our independent TB cohorts and confirm excellent specificity and sensitivity to distinguish active TB patients from those with LTBI. However, we identified dominance of IFN-inducible genes in these signatures and demonstrated enrichment of these signatures in published datasets of acute influenza infection^36,37^, which represents the immune response globally observed in viral infections. However, this signature was not apparent in bacterial pneumonia^36^, highlighting the apparent lack of IFN-inducible genes in the immunological response to bacterial pneumonia. However, it is clear that the immune response in TB has a dominant IFN-inducible gene signature resembling that of viral infections. It follows that such IFN-inducible signatures, whilst optimised for discriminating active TB patients and healthy individuals, with and without LTBI, with high sensitivity, may also detect other pathologies and/or infectious diseases that may exhibit a similar clinical presentation. It is clear that IFN-inducible genes are dominant discriminators of active TB from healthy LTBI, leading to preferential selection of this gene set to define an optimal signature. However, this dominance precludes consideration of most other gene sets and may also detect other diseases, such as viral infections. This view is supported by differences in the reported signatures of Kaforou et al.^25^ that were independently derived to discriminate active TB from LTBI, or active TB from other diseases. The 44-gene signature, derived using the latter approach, included more genes and exhibited greater diversity, when compared with the 27-gene signature for discriminating LTBI from TB. However, both signatures^25^ in addition to others^28,33,44,45^ still showed high specificity and sensitivity for influenza. These observations suggest that a trade-off exists between these two objectives, of achieving high sensitivity and high specificity, and that a single signature may not be optimal for both. The development of biomarkers for clinical practice is defined and optimised according to the clinical context for use. In a clinical context, the two objectives of a TB signature fulfil distinct requirements. A signature that discriminates active TB from LTBI is a useful screening tool for testing in healthy populations. Identification of an active TB signature when screening for LTBI can inform the need for further investigation. In contrast, a signature that discriminates active TB from other diseases would be applied for the investigation of unwell patients presenting with symptoms that suggest the possibility of TB, but may also be other infections including viral infections. While discrimination of TB from LTBI in screening programmes, and TB from other diseases in patients that are unwell, represent distinct clinical settings, the potential requirement of two different biomarkers adds complexity to models of implementation, particularly in a field setting. Furthermore, there is overlap between screening and clinical diagnostics as people attending for screening may present with an intercurrent illness (either symptomatic or asymptomatic) that is unrelated to TB. A single biomarker that is able to achieve reliable discrimination of TB from both LTBI and other conditions will have greater utility in clinical practice. A trade-off in sensitivity to distinguish active TB from LTBI may result from losing the IFN-inducible genes that are highly induced by viral infections. However, it is possible that this knowledge will allow the retention of such genes which distinguish active TB from LTBI with high sensitivity, by also including additional genes that are detected in response to viral infections and not TB, as an additional discriminatory approach to maintaining high sensitivity and specificity for TB against LTBI.

We developed the WGCNA derived modular signature for active TB across our cohorts and determined consistency of the signature in our cohorts and those from other published datasets. When taking into consideration all 23 modules, the signature in active TB was distinct from both viral and bacterial infections. In keeping with our earlier findings of an IFN-inducible signature of active TB^9^, we here also demonstrate an IFN-inducible gene signature, in both the active TB and LTBI outliers. The IFN-inducible signature, however, is now distributed across three different modules; two over-abundant modules; the yellow module that includes *BATF2, AIM2, FCGR1A and B*, and a number of *GBPs;* the Lightgreen module, which we show is also strongly over-abundant in influenza infection, includes many *IFITs, ISGs and OASs*, very reminiscent of type-I IFN inducible genes induced during viral infections. In contrast, the Tan module, which includes *IFNG* and *TBX21*, is significantly under-abundant in TB and some LTBI outliers, in keeping with the reported down-regulation of *IFNG* expression and signalling by high levels of type I IFN, contributing to the pathogenesis of TB^15^. This could also represent the reduced number of CD4^+^ and CD8^+^ T cells that we observed in the blood of active TB patients as we have previously discussed^9^. We also observed that this IFNG module was under-abundant in those contacts who progressed to TB (7 out of 9), whereas few of the IGRA^+ve^ (3 out of 16) and IGRA^−ve^ (2 out of 15), showed an under-abundance of this module. This supports the hypothesis that the ratio of type I IFN versus IFNG inducible genes, may be critical in determining protection or progression to TB disease.

To tackle the challenge of developing a high performing TB signature, we explored the modular tool for systematic gene reduction into biologically meaningful modules that together represent the entire transcriptome. Furthermore, we additionally identified gene clusters that were differentially expressed in TB but not influenza from within the modules of IFN signalling. This was an important observation as the opportunity to select specific genes from these dominant modules offered scope to improve the sensitivity of the signature and discrimination of TB from LTBI and other diseases. Based on these findings we developed and evaluated a two-step approach for targeted gene selection to derive a TB signature. Modules perturbed in TB were first interrogated to establish a gene set comprising genes that are differentially expressed in TB compared with other diseases. *A priori* gene selection in this way provided a gene set with high TB specificity against other diseases. In the second step, traditional gene reduction methodology was applied to separate TB from LTBI using this gene set. As a proof of principle, we developed a 20-gene signature using this approach that was diverse in its modular representation, incorporating genes from 6 modules. We demonstrate here that this 20-gene signature has robust sensitivity and specificity for discriminating active TB from LTBI in our cohorts, in addition to across a number of different published cohorts^10,25,28^. Our 20-gene signature also showed discrimination of TB from other diseases, similarly to the 44-gene signature of Kaforou et al.^25^. However, our 20-gene signature did not detect influenza from healthy controls, in contrast to all the other reported TB signatures^25,28,33,44,45^, which not only detected TB at high specific and sensitivity against LTBI, but additionally showed a high specificity and sensitivity for influenza versus controls. Therefore, the development of our 20-gene signature provides a novel approach to discriminate TB from LTBI, whilst not detecting viral infections, here exemplified by influenza, offering scope for further refinement in further translational clinical studies.

Heterogeneity of LTBI was suggested in our previous study^9^ with the identification of an outlier group after clustering. In the present study, we identified a similar proportion of LTBI outliers in the new Leicester cohort. We demonstrated enrichment scores using the published signatures of Zak et al.^33^ and Kaforou et al.^25^, dominated by IFN-inducible genes, that were higher in outliers compared with other LTBI in both the Berry and Leicester cohorts, and overlapped with scores obtained in active TB. These observations suggested LTBI outliers are characterised by an overabundance of IFN-inducible genes, a view that was corroborated in their modular signatures, together with identification of seventy selectively upregulated genes, common to both the Berry and Leicester LTBI outliers, which mapped to IFN signalling pathways. The clinical significance of these observations remains unclear, however the recent study of Zak et al.^33^ suggests expression of a TB-like signature, characterised by enrichment of IFN-inducible genes which we show from our analysis, may indicate either subclinical disease or increased risk of progression to TB within a few months, although this may be confounded by viral infections.

We utilised the modular signature for deeper characterisation of heterogeneity in recent TB contacts and identified instances of similarity with TB in a few of the IGRA^−ve^ (2) and IGRA^+ve^ (4) individuals, representing IFN and other signalling pathways. There were higher perturbations in the modular signature in IGRA^+ve^ individuals. Our observations of low modular activity for the IGRA^−ve^ cohort is consistent with the absence of LTBI and likely to reflect a robust finding. In contrast, enrichment scores of the published signatures we tested indicated considerably more overlap of IGRA^−ve^ subjects with the IGRA^+ve^ group, again indicating impaired specificity of these signatures. Other modular changes, discordant with TB, appeared to be driven by differences in the pattern of perturbation in modules than those representing IFN signalling pathways. The majority of contacts who developed TB had a modular signature (6 out of 9) comparable to that of active TB patients, observable before a diagnosis was made.

In keeping with the global modular activity, we observed evidence of dynamic change in the reduced 20-gene signature derived from the modules, of some TB contacts that can be categorised into three patterns of longitudinal expression that may reflect early immunological events following TB exposure. We suggest the absence of a signature at any time point may indicate the absence of infection being acquired. This pattern was seen in 67% of our IGRA^−ve^ cohort and 38% of our IGRA^+ve^ cohort. A transient signature may indicate an infection that was acquired but has either been controlled or cleared. In this context, the observation that 26% of our IGRA^−ve^ cohort and in 50% of our IGRA^+ve^ cohort demonstrated this pattern suggests that the blood transcriptional signature represents immune responses that may precede priming and activation of IFN - y producing CD4 T-cells. Finally, subjects with an evolving and persistent modular TB signature may represent subjects that have acquired an infection requiring active control to maintain latency. This pattern was seen in 7% of IGRA^−ve^ subjects and 12% of IGRA^+ve^. These observations require validation in larger longitudinal cohorts, but do suggest that the blood transcriptome may offer a more sensitive approach to characterising the state of latent infection following TB exposure, with implications for better stratification of prospective TB risk.

For our cohort of 9 subjects identified with TB during prospective observation, a high or rising 20-gene signature score was observed in most. This was most apparent in the subjects defined as true progressors, who had no signature at baseline. Our study was limited by small numbers and the identification of TB within a short period of prospective observation, suggesting that incipient TB is likely to have been present at the time of baseline assessment in a proportion of cases. We are therefore presently unable to comment on the dynamic properties of this response or determine accurately the interval between the signature becoming detectable and manifestation of active TB. It is notable also that three subjects did not express a signature at any time point and yet went on to be diagnosed with TB. One of these was on anti-bacterial drugs which have known immunosuppressive properties, which could have diminished the signature. Additionally, interrogating the modules for these subjects indicates a weak transcriptional response that may suggest pathogen induced host immunomodulation, which is well recognised in active TB^16,46^. We are unable to determine whether a delayed transcriptional response would have developed over the natural time-course of infection as our rigorous protocol of frequent surveillance identified active disease at the earliest stage, however it is apparent that heterogeneity of the host immune response and its association with the state of *M.tuberculosis* requires further investigation.

A robust, objective definition of LTBI is unavailable. Patients with IGRA positivity have a heterogeneous risk of developing TB and secondly, a proportion of patients that are IGRA negative at screening proceed to develop TB in the future. It therefore follows that an IGRA is not a reliable gold standard to determine the validity of new biomarkers. In this respect, our observations of a TB like modular signature being expressed in both IGRA positive and IGRA negative contacts of TB is not surprising. The difference between the groups in the proportion of subjects expressing the signature (19% in IGRA^+ve^ vs 7% in IGRA^−ve^ is comparable with the relative risk of TB according to IGRA status (Incident rate ratio for TB 2.11,^57^). This provides support for the validity of our observations and supports developing a transcriptional biomarker for defining LTBI.

Here in summary, we have validated whole blood transcriptomic findings, previously identified by microarray by RNA-sequencing of our previously published TB cohorts and a new cohort from a low-TB-incidence setting. We further developed an advanced modular signature of active TB, and validated it in our new cohort and a number of TB cohorts published by other groups. Using this modular signature, we obtained a reduced TB-specific 20-gene signature which showed very high specificity and sensitivity in individuals with active TB against those with LTBI and other diseases. Moreover, this signature did not detect influenza, representative of many viral infections that share a strong IFN-inducible signature, providing a proof-of-principle for the development of transcriptional biomarkers for TB as diagnostics, with the aim of obtaining the highest sensitivity, whilst maintaining specificity against LTBI and other diseases. Our findings highlight the need to consider additional approaches to develop transcriptional biomarkers of the highest sensitivity and specificity for distinguishing active TB from LTBI and other diseases. The reduced gene signatures for discriminating active TB from LTBI and other infections, also demonstrated important clinical outcomes and heterogeneity in LTBI. Our improved approach for the development of diagnostic biomarkers consisting of reduced gene-sets, is broadly applicable across diverse infectious and inflammatory diseases.

## Methods

### Study cohorts for analysis

Cohorts analysed in Berry et al. 2010^9^ using microarrays were subjected to RNA-Seq and analysed as part of this study. Test and validation sets, termed Berry London and Berry South Africa sets, respectively, based on the geographical location of patient recruitment, were retained for RNA-seq analysis in this study (**Supplementary Figure 1a**).

An independent cohort was recruited (between 09/2015 and 09/2016) at the Glenfield Hospital, University Hospitals of Leicester NHS Trust, Leicester, UK. The cohort consisted of active TB patients (n=53) and recent close contacts (n=108). Patients who were pregnant, immunosuppressed, had previous TB or previous treatment for LTBI were excluded from this study. All participants had routine HIV testing and patients with a positive result were excluded. Patients with active TB were confirmed by laboratory isolation of *M. tuberculosis* on culture of a respiratory specimen (sputum or bronchoalveloar lavage) with sensitivity testing performed by the Public Health Laboratory Birmingham, Heart of England NHS Foundation Trust, Birmingham, UK. All recent close contacts were IGRA tested using the QuantiFERON Gold In-Tube Assay (Qiagen) and were subsequently categorised as either IGRA negative (n=50) or IGRA positive (n=49). All participants were prospectively enrolled and sampled before the initiation of any anti-mycobacterial treatment. A subset of subjects recruited initially as close contacts were identified with active TB during longitudinal assessment (n=9), based on microbiological confirmation of *M. tuberculosis* by culture or positive Xpert MTB/RIF (Cepheid). (**Supplementary Tables 1** and **5; Figure 7a**). The Research Ethics Committee (REC) for East Midlands - Nottingham 1, Nottingham, UK (REC 15/EM/0109) approved the study. All participants were older than 16 years and gave written informed consent.

### RNA extraction and cDNA library preparation for RNA-Seq

3 ml whole blood were collected by venepuncture into Tempus™ blood RNA tubes (Fisher Scientific UK Ltd), tubes were mixed vigorously immediately after collection, and then stored in a −80°C freezer prior to use. Total RNA was isolated from 1 ml whole blood using the MagMAX™ for Stabilized Blood Tubes RNA Isolation Kit (Applied Biosystems/Thermo Fisher Scientific) according to the manufacturer’s instructions. Globin RNA was depleted from total RNA (1.5-2 μg) using the human GLOBINclear kit (Thermo Fisher Scientific) according to manufacturer’s instructions. RNA yield of total and globin-reduced RNA was assessed using a NanoDrop™ 8000 spectrophotometer (Thermo Fisher Scientific). Quality and integrity of total and globin-reduced RNA were assessed with the HT RNA Assay reagent kit (Perkin Elmer) using a LabChip GX bioanalyser (Caliper Life Sciences/Perkin Elmer) and assigned an RNA Quality Score (RQS). Samples (200 ng) with an RQS > 6 were used to prepare a cDNA library using the TruSeq Stranded mRNA HT Library Preparation Kit (Illumina). The tagged libraries were sized and quantitated in duplicate (Agilent TapeStation system), using D1000 ScreenTape and reagents (Agilent), normalised, pooled and then clustered using the HiSeq® 3000/4000 PE Cluster Kit (Illumina). The libraries were imaged and sequenced on an Illumina HiSeq 4000 sequencer using the HiSeq® 3000/4000 SBS kit (Illumina) at a minimum of 25 million paired end reads (75 bp) per sample.

### RNA-seq data analysis

Raw paired-end RNA-seq data obtained for Berry London, Berry South Africa and Leicester cohorts was processed separately and subjected to quality control using FastQC (Babraham Bioinformatics) and MultiQC^58^. Trimmomatic^59^ v0.36 was used to remove adapters and filter raw reads below the 36 bases long and leading and trailing bases below quality 25. Filtered reads were aligned to the *Homo sapiens* genome Ensembl GRCh38 (release 86) using HISAT2^60^ v2.0.4 with default settings and RF rna-strandedness including unpaired read reads resulting from Trimmomatic. Mapped and aligned reads were quantified to obtain gene-level counts using HtSeq^61^ v0.6.1 with default settings and reverse strandedness. Raw counts were processed using the *bioconductor* package edgeR^62^ v3.14.0 in R. Genes expressed with counts per million (CPM) >2 in at least 5 samples were considered and normalised using trimmed mean of M-values (TMM) to remove library-specific artefacts. Only protein coding genes were considered for subsequent analyses. Differentially abundant genes were calculated using likelihood ratio tests in edgeR by fitting generalized linear models to the non-normally distributed RNA-seq data. Genes with log2 fold change >1 or <−1 and false discovery rate (FDR) p-value < 0.05 corrected for multiple testing using the Benjamini-Hochberg (BH) method^63^ were considered significant. For subsequent analysis, voom transformation was applied to RNA-seq count data to obtain normalized expression values on the log2 scale. For Berry Combined dataset, raw counts from Berry London and South Africa cohorts were combined as one dataset and processed in edgeR as described above and batch effects were removed from log2 expression values using surrogate variable analysis (sva) using the *bioconductor* package sva^64^ in R. RNA-seq data obtained from Zak et al. 2016^33^ in the SRA format were converted to fastq files using the SRA toolkit and processed as above.

### Microarray data analysis

External microarray datasets retrieved from GEO as non-normalized matrices were processed in GeneSpring GX v14.8 (Agilent Technologies). Flags were used to filter out probe sets that did not result in a ‘present’ call in at least 10% of the samples, with the ‘present’ lower cut-off of 0.8. Signal values were then set to a threshold level of 10, log2 transformed, and per-chip normalised using 75^th^ percentile shift algorithm. Next per-gene normalisation was applied by dividing each messenger RNA transcript by the median intensity of all the samples. The training, test and validation sets in Bloom et al. 2013^10^ were combined and batch effects were removed using sva^64^. In Kaforou et al. 2013^25^, HIV+/− groups were combined and analysed as one dataset. In all datasets, multiple probes mapping to the same gene were removed and the probe with the highest inter-quartile range across all samples was retained, to match with the RNA-seq data. Differentially expressed genes were identified using the *bioconductor* package *limma*^65^ in R and only genes with FDR p-value < 0.05 corrected for multiple testing using the BH method^63^ were considered significant.

### Gene signature enrichment analysis

Enrichment of TB gene signatures was carried out on a per sample basis using single sample Gene Set Enrichment Analysis (ssGSEA)^35^ using the *bioconductor* package *gsva*^66^ in R. Enrichment scores were obtained similar to those from Gene Set Enrichment Analysis (GSEA) but based on absolute expression rather than differential expression^35^, to quantify the degree to which a gene set is over-represented in a particular sample.

### Weighted gene co-expression network analysis

Modular analysis was performed using the WGCNA package in R. Modules were constructed using the Berry Combined dataset (combined Berry London and South Africa sets) using 5,000 genes with highest covariance across all samples using log2 RNA-seq expression values. A signed weighted correlation matrix containing pairwise Pearson correlations between all genes across all samples was computed using a soft threshold of β = 14 to reach a scale-free topology. Using this adjacency matrix, the Topological Overlap Measure (TOM) was calculated, which measures the network interconnectedness and used as input to group highly correlated genes together using average linkage hierarchical clustering. The WGCNA dynamic hybrid tree-cut algorithm^67^ was used to detect network modules of co-expressed genes, with a minimum module size of 20. All modules were assigned a colour arbitrarily and annotated using Ingenuity Pathway Analysis (IPA) (QIAGEN Bioinformatics) and Literature Lab (Acumenta Biotech, Massachusetts, USA). Literature Lab mines the PubMed literature and identifies significant associations in 20 MeSH (Medical Subject Headings) domains, including pathways, diseases and cell biology. Significantly enriched canonical pathways from IPA (p-value<0.05), and strongly associated terms from Literature Lab were obtained. Modules were assigned annotation terms based on pathways and processes that showed corroboration between both tools (**Supplementary Table 4**). Representative terms were then selected and assigned to modules (**Figure 4**). For each module, module eigengene (ME) values were calculated, which represent the first principal component of a given module and summarize the gene abundance profile in that module. For each module, top 50 hub genes with high intramodular connectivity and a minimum correlation of 0.75 were calculated and exported into Cytoscape v3.4.0 to create interaction networks.

### WGCNA module enrichment analysis

Fold enrichment for the WGCNA modules was calculated using quantitative set analysis for gene expression (QuSAGE)^68^ using the *bioconductor* package *qusage* in R, to identify the modules of genes over- or under-expressed in a dataset compared to a control group. Linear mixed models were incorporated in the analysis using QGen algorithm in QuSAGE, and patients in datasets with repeated measures were modelled as random effects. Only modules with FDR p-value < 0.05 were considered significant. To test the modules in microarray datasets, only those modules with a >70% match in genes symbols was present in the microarray dataset. To obtain a modular profile of a disease group, single sample enrichment scores were calculated using ssGSEA and the average enrichment score of the control group was subtracted from the average enrichment score of the disease group. To obtain a modular profile on a single sample basis, average enrichment score of the control group was subtracted from the enrichment score of the sample.

### Class prediction

In order to develop a TB-specific gene signature, only genes significantly differentially expressed in Berry London set and not in other flu cohorts were considered, from only those modules that were perturbed in TB (a module was considered perturbed in TB if it followed a similar profile (up or down compared to control) in at least 4 of the 5 TB datasets (Berry London, Berry South Africa, Leicester cohort, Kaforou et al. 2013^25^ and Zak et al. 2016^33^), and given that for the 5^th^ dataset the module did not reach significance when compared to control). These genes were then reduced using the *Boruta*^42^ package in R. Boruta is a feature selection wrapper algorithm based on random forest and is particularly useful in biomedical applications as it captures features by incorporating the outcome variable. Next, the features identified as predictive using Boruta were ranked using the GINI score in random forest and the top 20 genes were selected. For classifying patients as active TB or latent TB, the random forest algorithm was used in *caret*^69^ package in R, using LOOCV over 1,000 iterations. Each of the TB datasets was randomly split into training (70%) and test (30%) sets to classify patients as active TB or latent TB. For the Zhai et al. 2015^37^, the Influenza A group at Day 0 was randomly split into training (70%) and test (30%) sets to classify patients as infected with Influenza A or healthy controls.

### Modified Disease Risk Score

To test the TB-specific 20-gene signature, a modified version of the Disease Risk Score (DRS) established by Kaforou et al. 2013^25^ was used. Briefly, the DRS is obtained from normalized data in a non-log space, by adding the total intensity of up-regulated transcripts and subtracting the total intensity of down-regulated transcripts from a gene signature. In this study, normalized CPM values were used for the RNA-seq data and non-log normalized expression values were used for microarray data. As part of the modification of the DRS, the absolute values of the total intensity of up-regulated transcripts and total intensity of down-regulated transcripts were added to obtain a composite score.

### Deconvolution analysis

Deconvolution analysis for quantification of relative levels of distinct cell types on a per sample basis was carried out using CIBERSORT^70^. CIBERSORT estimates relative subsets of RNA transcripts using linear support vector regression. Cell signatures for 22 cell types were obtained using the LM22 database from CIBERSORT and grouped into 11 representative cell types. Fractions of cell types were compared across different groups using One-way ANOVA, and p-value < 0.05 was considered significant.

### Data availability

#### Novel data newly generated in this study

Sequence data generated for our study that support the findings have been deposited in NCBI GEO database with the primary accession code <u>GSE107995</u>, and in BioProject with the primary accession code <u>PRJNA422124</u>. Additional data which we have generated that support the findings of our study are available upon request.

#### Data we accessed from the literature and GEO for our study

The TB datasets from the literature that have been used in this study as comparators are available in GEO with the primary accession codes <u>GSE37250, GSE79362</u> also in BioProject with the primary accession code <u>PRJNA315611</u>, and in SRA with the primary accession code <u>SRP071965, GSE20346, GSE68310, GSE42026, GSE60244</u> and <u>GSE42834</u>.

## Acknowledgements

We acknowledge the Francis Crick Advanced Sequencing Facility, and Bioinformatics and Biostatistics Science Technology Platforms for their contribution to our sequencing processing. We acknowledge the NIHR Leicester Biomedical Research Centre for their support of the study at Leicester. The views expressed are those of the author(s) and not necessarily those of the NHS the NIHR or the Department of Health. We thank the patients for their participation. We thank Asmaà Fritah-Lafont for help in co-ordinating the meetings regarding the study. We thank Dr. Lúcia Moreira-Teixeira for reviewing the manuscript and for valuable discussion. AOG, CMG and AS were funded by The Francis Crick Institute, (Crick 10126; Crick 10468), which receives its core funding from Cancer Research UK, the U.K. Medical Research Council, and the Wellcome Trust; and the sequencing project by the BIOASTER Microbiology Technology Institute, Lyon, France; Medical Diagnostic Discovery Department, bioMérieux SA, Marcy l’Etoile, France; and funded in part by Illumina Inc., San Diego, CA, USA. RV and JL, University of Leicester, were funded by BIOASTER Microbiology Technology Institute, Lyon, France. This work has received, through BIOASTER investment, funding from the French Government through the Investissement d’Avenir program (Grant NO. ANR-10-AIRT-03). RJW was supported by The Francis Crick Institute, (Crick 10128), which receives its core funding from Cancer Research UK, the U.K. Medical Research Council, and Wellcome; by Wellcome (104803; 203135); MRC South Africa under strategic health innovation partnerships; and NIH 019 AI 111276.

## Author contributions

AOG and PH co-led the whole study; AOG, MPRB, PH, RV, GW, MRo designed the study; RV and JL recruited TB, LTBI and contacts to the study for the Leicester cohort; CMG led and performed the RNA-Seq sample and raw data generation. RV and CMG helped to co-ordinate logistics of the study; RJW, PLei, PLec, KK gave feedback and concrete discussion during the study; TT and MRi contributed towards feedback on bioinformatics analysis; AS led and performed all the bioinformatics analysis;. AOG, AS, RV and PH wrote the manuscript and RJW gave substantial input; all co-authors have read, reviewed and approved the paper.

ORCID IDs: 0000-0001-9845-6134 (AOG); 0000-0002-6941-3618 (AS)

